# Neurotrophic Factor-α1/carboxypeptidase E regulates critical protein networks to rescue neurodegeneration, defective synaptogenesis and impaired autophagy in Alzheimer’s Disease mice

**DOI:** 10.1101/2025.06.04.657876

**Authors:** Lan Xiao, Pranav Sharma, Xuyu Yang, Daniel Abebe, Y. Peng Loh

**Author notes:** **Corresponding author :** Dr. Y. Peng Loh, Section on Cellular Neurobiology, 49, Convent Drive, Bldg. 49, Rm 6A-10, NICHD, NIH, Bethesda, MD 20892, USA.

## Abstract

**Background:** The global aging population is increasingly inflicted with Alzheimer’s disease (AD), but a cure is still unavailable. Neurotrophic Factor-α1/carboxypeptidase E (NF-α1/CPE) gene therapy has been shown to prevent and reverse memory loss and pathology AD mouse models However, the mechanisms of action of NF-α1/CPE are not fully understood. We investigated if a non-enzymatic form of NF-α1/CPE-E342Q is efficient in reversing AD pathology and carried out a proteomic study to uncover the mechanisms of action of NF-α1/CPE in AD mice.

**Methods:** AAV-human NF-α1/CPE and a non-enzymatic form, NF-α1/CPE -E342Q were delivered into hippocampus of 3xTg-AD mice and effects on cognitive function, neurodegeneration, synaptogenesis and autophagy were investigated. A quantitative proteomic analysis of hippocampus of 3xTg-AD mice with and without AAV-NF-α1/CPE treatment was carried out.

**Results:** Hippocampal delivery of AAV-NF-α1/CPE-E342Q prevented memory loss, neurodegeneration and increase in activated microglia in 3xTg-AD mice, indicating its action is independent of its enzymatic activity. Quantitative proteomic analysis of hippocampus of 3xTg-AD mice that underwent NF-α1/CPE gene therapy revealed differential expression of >2000 proteins involving many metabolic pathways. Of these, two new proteins down-regulated by NF-α1/CPE: Nexin4 (SNX4) and Trim28 which increase Aβ production and tau levels, respectively were identified. Western blot analysis verified that they were reduced in AAV-NF-α1/CPE treated 3xTg-AD mice compared to untreated mice. Our proteomic analysis indicated synaptic organization as top signaling pathway altered as a response to CPE expression. Synaptic markers PSD95 and Synapsin1 were decreased in 3xTg-AD mice and were restored with AAV-NF-α1/CPE treatment. Proteomic analysis hypothesized involvement of autophagic signaling pathway. Indeed, multiple proteins known to be markers of autophagy were down-regulated in 3xTg-AD mice, accounting for impaired autophagy. Expression of these proteins were upregulated in 3xTg-AD mice with NF-α1/CPE gene therapy, thereby reversing autophagic impairment.

**Conclusions:** This study uncovered vast actions of NF-α1/CPE in restoring expression of networks of critical proteins including those necessary for maintaining neuronal survival, synaptogenesis and autophagy, while down-regulating many proteins that promote tau and Aβ accumulation to reverse memory loss and AD pathology in 3xTg-AD mice. AAV-NF-α1/CPE gene therapy uniquely targets many metabolic levels, offering a promising holistic approach for AD treatment.

**Graphic abstract:** 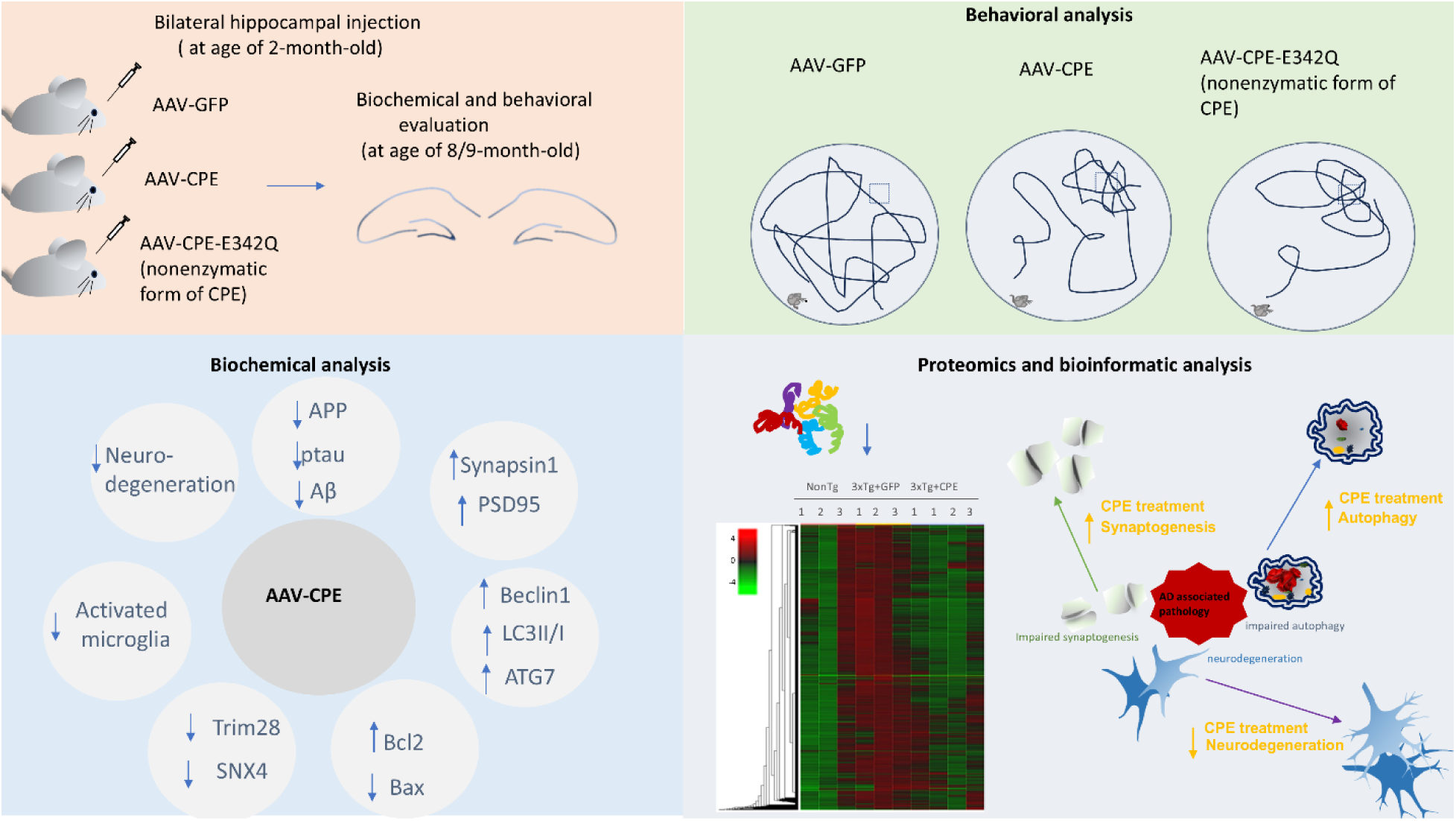

## Introduction

Alzheimer’s Disease (AD) is one of the most prevalent neurological disorders that lead to dementia worldwide. With an increasing ageing population globally, the number of patients diagnosed with AD is expected to reach 152 million by 2050 [1]. Both environmental and genetic components contribute to the pathogenesis of AD. Majority of patients demonstrated late-onset symptoms after age of 65, while approximately 5% of patients developed early-onset symptoms before age of 65 due to genetic predisposition such as for *Amyloid precursor protein,(APP),* and *presenilins 1*,2 [2]. Most AD patients demonstrate progressive memory loss, impaired visuospatial and executive function [3]. Pathologically, AD is characterized by severe neurodegeneration, abnormal aggregation of β-amyloid extracellularly and hyperphosphorylated tau intracellularly in the central nervous system. A variety of altered cellular activities are also closely related with the pathogenesis of AD. Autophagy plays a critical role in maintaining intracellular protein homeostasis. In the major autophagy-lysosome pathway, misfolded proteins or damaged organelles are encapsulated within autophagosomes and then transported to lysosomes for degradation. A large number of studies have reported that autophagy dysfunction is highly associated with accretion of amyloid and neurodegeneration [4]. In the brains of AD patients and PS1/APP mouse models, autophagosomes accumulated in the dystrophic neurites and contributed to the aggregation of beta amyloid [5]. On the other hand, reduction of synapses and synaptogenesis are also correlated with declined cognition and exacerbation of AD [6, 7]. Specially, pathological tau is linked with synaptic loss and synaptic dysfunction [8]. Although, AD has been known for more than a hundred years since first described in 1906 [9, 10], the regulation of the deficits remains poorly understood.

Neurotrophic Factor-α1/Carboxypeptidase E (NF-α1/CPE), which has been shown to exert neuroprotective effects both in *in vitro* and *in vivo* studies [11, 12] has recently been demonstrated to prevent and reverse memory loss and AD pathology in three different mouse models [13]. CPE-KO mice displayed hippocampal CA3 neurodegeneration and impaired learning and memory activity [14]. Mice carrying mutated CPE genes also demonstrated abnormal hippocampal function and deficits in cognition[14, 15]. In humans, association between CPE mutation and cognitive impairment and learning disability has been reported in several clinical cases [16, 17]. Interestingly, studies have indicated that although CPE plays a critical role in processing proneuropeptide, its neuroprotective activity is independent of its enzymatic activity. This is supported by studies showing that transgenic mice carrying a nonenzymatic form of CPE-E342Q displayed intact hippocampal structure and memory and learning, in comparison with CPE-KO mice[12].

Studies in human neurons support the involvement of CPE in neuroprotection. CPE is mainly distributed in neurons, astrocytes and glia in the cortex of humans. In the cortex of AD patients, CPE was observed accumulating in dystrophic neurites around amyloid [18]. Another study showed that NF-α1/CPE was co-localized with the serotonin receptor HTR1E in human hippocampal neurons. Secreted NF-α1/CPE interacts with HTR1E to activate ERK-CREB signaling pathway and upregulates Bcl2 signaling, which convergently enhances neuronal survival [19].

Our previous studies have shown that hippocampal delivery of AAV-NF-α1/CPE (mouse) in 3xTg -AD mouse model prevented cognitive decline, neurodegeneration, and amyloid and tau pathology. To further elucidate the mechanisms by which NF-α1/CPE rescues AD pathology, in this study, we have injected AAV-human NF-α1/CPE or a non-enzymatic form of human NF-α1/CPE (E342Q) in the hippocampus of 3xTg-AD mice and carried out an extensive proteomic analysis to identify critical proteins that are modulated by NF-α1/CPE to rescue cognitive dysfunction, neurodegeneration and impaired synaptogenesis and autophagy, characteristic of AD. The AAV-NF-α1/CPE and NF-α1/CPE-E342Q treatment rescued memory deficit and neurodegeneration observed in 9-month-old 3xTg-AD mice. Proteomic analysis revealed that NF-α1/CPE regulates expression of many critical proteins to mitigate neurodegeneration, impaired autophagy and synaptogenesis observed in the 3xTg-AD mice.

## Material and methods

### Animals

The mouse strain used for this research project, B6;129-Tg(APPSwe,tauP301L)1Lfa Psen1tm1Mpm/Mmjax, RRID:MMRRC_034830-JAX, was obtained from the Mutant Mouse Resource and Research Center (MMRRC) at The Jackson Laboratory, an NIH-funded strain repository, and was donated to the MMRRC by Frank Laferla, Ph.D., University of California, Irvine; Mark P. Mattson, Ph.D., Johns Hopkins University, School of Medicine.” [20–23].

All mice were housed at NIH animal facility with free access to food and water ad libitum and controlled humidity (45%) and temperature (22°C) under a 12h light/dark cycle. At age of ∼2 months, 3xTg-AD and nonTg mice were randomly selected and divided into 4 groups: nonTg+GFP, 3xTg+GFP, 3xTg+ NF-α1/CPE and 3xTg+ NF-α1/CPE-E342Q to receive bilateral hippocampal stereotaxic injections. At age of ∼8 months, brain tissues were collected for biochemical and immunohistochemical studies after behavioral tests.

### Viral vectors

AAV1/2-GFP and AAV1/2-CPE (chimeric serotype) viral constructs were purchased from Vector Biolabs (Philadelphia, PA). AAV1 and AAV2 transduce neurons, reactive microglia and astrocytes. GFP and CPE expressions in these AAV constructs are driven by the CMV promoter.

### Stereotaxic injection

Stereotaxic injection was conducted as previously described [13]. AAV viruses expressing GFP or human NF-α1/CPE or NF-α1/CPE-E342Q or were bilaterally injected into the hippocampus (total 1×10^10^ VP,1ul on each side of hippocampus) according to the coordinates AP :-1.94mm, L: ± 1.0mm, V:-1.3mm.

### Electronic microscopy (EM)

Mice were transcardially perfused, fixed with 4% PFA with 2.5% glutaraldehyde made in PBS buffer, pH 7.4. Brains were removed and left to post-fix overnight in the same fixative at 4° C. After primary fixation and dissection, samples were cut at 100um on a Vibratome ( LEICA VT1000 S). Next, regions of interest were selected, cut and inserted into mPrep tissue capsules and loaded onto an mPrep ASP-2000 Automated Biological Specimen Preparation Processor (Microscopy Innovations, LLC, Marshfield, WI.) which automates all processing steps which included the following steps: post-fixation in 2% osmium tetroxide, en-bloc in 2% uranyl acetate (aqueous), dehydrated in a graded ethanol series followed by further dehydration in100% acetone and finally infiltrated and embedded in Embed 812 epoxy resin (Electron Microscopy Sciences Hatfield, PA.). Embedded samples were polymerized in an oven set at 60°C. Samples were then ultra-thin sectioned (90nm) on a Leica EM UC7 Ultramicrotome. Thin sections were picked up and placed on 200 mesh copper grids and post-stained with UranyLess (Uranyl Acetate substitute, Electron Microscopy Sciences Hatfield, PA.) and lead citrate. Imaging was performed on a JEOL-1400 Transmission Electron Microscope operating at 80kV and an AMT BioSprint-29 camera.

### Behavioral studies

Male nonTg and 3xTg-AD mice at 2 months of age were injected with either AAV-GFP or AAV-NF-α1/CPE or NF-α1/CPE-E342Q in the hippocampus and then tested for open field and Morris water maze at ∼8 months of age.

#### Open field test

To evaluate the locomotor activity, mice were tested in open field apparatus for 1 h, and the distance and speed mice traveled were evaluated by ANY-maze (ANY-maze, Wood Dale, IL).

#### Morris Water Maze test

Morris water maze was conducted as previously described [24]. Morris water maze consists of two sessions: a hidden platform training session (day1-5) and a probe test session (day 6). The test was performed in a circular pool full of water and nontoxic white paint. On day 1-5, the hidden platform was positioned in the same spot and mice were allowed to search for the platform for 1 min. There were four trials each day and mice were placed in a new quadrant of the pool in each trial. If mice failed to find the hidden platform, they were guided to the platform and allowed to sit on it for 30 seconds. On day 6, the hidden platform was removed, and mice were allowed to explore the pool for 1min. The time in each quadrant was recorded and analyzed by ANY-maze(ANY-maze, Wood Dale, IL).

### Aβ40 and Aβ42 ELISA

Mouse brain tissues were processed as previously described [25]. Hippocampal tissues were homogenized and centrifuged for 30min at 100,000g at 4 C. The supernatant which contains soluble proteins was collected and saved. The precipitate which contains the insoluble proteins was resuspended in 70% formic acid (Sigma-Aldrich, St. Louis, MO). The supernatant and precipitate were analyzed for soluble and insoluble Aβ40 and Aβ42 using an ELISA kit specific for human, following the manufacture’s protocol (Invitrogen, Waltham, MA).

### Western blot

Mouse brain tissues were prepared as previously described [24]. Hippocampal lysates were run on SDS-PAGE gels and transferred onto nitrocellulose membrane. The membrane was then incubated with primary antibodies (Supplementary Table 1) overnight followed by secondary fluorescent conjugated anti-mouse or rabbit antibodies. The protein expressions were visualized and quantified by the Odyssey infrared imaging system (LI-COR Inc, Lincoln, NE) and normalized to β-actin.

### Immunohistochemistry

Mouse brains were sectioned coronally at 25µm and then incubated with primary antibodies (Supplementary Table 2) and then followed with 1:1000 biotinylated (Vector, Newark, CA) or 1:500 fluorescence (Jackson Lab, Bar Harbor, ME) secondary antibodies. Images were scanned with an Olympus VS200 slide scanning system. For MAP2 and GFAP quantification, two random areas in CA1 region (167umx167µm) (four sections per animal, six animals per genotype) were selected. MAP2 intensity was measured by Image J and GFAP positive cells were counted. To quantify CD68 positive cells, numbers of positive cells and total cells in CA1 region were counted within an area (approximately 160μm x 320μm). 4 sections per mouse, 6 mice per genotype. Percentage of positive cells versus total cell numbers was calculated.

### Proteomic Study

#### Cell lysate preparation

Hippocampus were dissected from 3 groups of mice: nonTg+GFP, 3xTg+GFP, 3xTg+CPE in triplicates and cell lysate were prepared in RIPA buffer (Thermofisher, Waltham, MA) completed with Halt™ Protease and Phosphatase Inhibitor Cocktail (Thermo Scientific, Waltham, MA).

#### Protein digestion protocol

Protein concentration was determined using micro-BCA (Thermo Fisher) for each of the 9 cell lysates. Equal amount of protein for each group was precipitated using ammonium sulphate. The pellets were resuspended in 600ul of 8M urea made in 100mM Tris pH 8.0 by vortexing for 5-10 minutes. Tris(2-carboxyethyl)phosphine hydrochloride (TCEP) was added to the final concentration of 10 mM. The sample was then frozen overnight at -20°C to help solubilization of the proteins in urea solution. Next day, the solution was thawed and vortexed for another 5 minutes until the solution became clear. The Chloro-acetamide solution was added to the final concentration of 40 mM and vortexed for 5 minutes. Equal volume of 50mM Tris pH 8.0 was added to the sample to reduce the urea concentration to 4 M. Lys C was added in 1:500 ratio of LysC to protein content and incubated at 37°C in a rotating incubator for 4-6 hours. Equal volume of 50mM Tris pH 8.0 was added to the sample to reduce the urea concentration to 2M. Trypsin was added in 1:50 ratio of trypsin to protein and incubated overnight at 37°C. The solution was then acidified using TFA (0.5% TFA final concentration) and vortexed for 5 minutes followed by centrifugation at 14,000g for 5 min to obtain aqueous and organic phases. The lower aqueous phase was collected and desalted using 100 mg C18-StageTips as described by the manufacturer protocol. The peptide concentration of sample was measured using BCA after resuspension in iTRAQ dissolution buffer.

#### TMT labeling

TMT tags from Thermo Scientific™ TMT10™ 10plex was used for the labeling using protocol as described by manufacturer.

#### High pH fractionation

Pierce™ High pH Reversed-Phase Peptide Fractionation Kit (Pierce™ High pH Reversed-Phase Peptide Fractionation Kit Catalog number: 84868) was used to fractionate labeled peptides using protocol as described by the manufacturer kit.

#### LC-MS-MS

Each fraction was analyzed by ultra-high-pressure liquid chromatography (UPLC) coupled with tandem mass spectroscopy (LC-MS/MS) using nano-spray ionization. The nano-spray ionization experiments were performed using an Orbitrap fusion Lumos hybrid mass spectrometer (Thermo) interfaced with nanoscale reversed-phase UPLC (Thermo Dionex UltiMate™ 3000 RSLC nano System) using a 25 cm, 75-micron ID glass capillary packed with 1.7-µm C18 (130) BEH^TM^ beads (Waters corporation). Peptides were eluted from the C18 column into the mass spectrometer using a linear gradient (5–80%) of ACN (Acetonitrile) at a flow rate of 375 μl/min for 180 min. The buffers used to create the ACN gradient were Buffer A (98% H_2_O, 2% ACN, 0.1% formic acid) and Buffer B (100% ACN, 0.1% formic acid). Mass spectrometer parameters are as follows; an MS1 survey scan using the orbitrap detector (mass range (m/z): 400-1500 (using quadrupole isolation), 60000 resolution setting, spray voltage of 2200 V, Ion transfer tube temperature of 275 C, AGC target of 400000, and maximum injection time of 50 ms) was followed by data dependent scans (top speed for most intense ions, with charge state set to only include +2-5 ions, and 5 second exclusion time, while selecting ions with minimal intensities of 50000 in which the collision event was carried out in the high energy collision cell (HCD Collision Energy of 38%) and the first quadrupole isolation window was set at 0.7 (m/z). The fragment masses were analyzed in the Orbi-trap mass analyzer with a mass resolution setting of 15000 (with ion trap scan rate of turbo, first mass m/z was 100, AGC Target 20000 and maximum injection time of 22ms). Protein identification and quantification were carried out using Peaks Studio X (Bioinformatics solutions Inc).

#### Differential expression analysis

Proteomic analyses were performed on mouse brain samples, with protein intensities normalized by applying log2 transformation to reduce skewness and median normalization to account for variations in sample loading and global batch intensity differences. Principal component analysis (PCA) implemented with the R package *prcomp*, was used to evaluate the influence of covariates such as batch, sex, age, and genotype, ensuring that biological signals were not confounded by confounding variables. Differential expression (DE) analysis was performed using a moderated *t*-test from the *MKmisc* package, which estimates the variance by borrowing information across all proteins, providing increased statistical power compared to traditional *t*-tests. To control for false discoveries in multiple testing, the *p* values were adjusted to FDR using the Benjamini-Hochberg procedure.

#### Pathway enrichment and protein–protein interaction network analysis

Pathway enrichment of DEPs was performed by using STRING software. Pathway enrichment specifically emphasized GO categories relevant to neurodegenerative processes, informed by prior knowledge of AD pathogenesis. Results were filtered by false discovery rate (FDR) (below 0.01) to identify DEP-associated pathways with high confidence.

## Statistical analysis

Data are representative of at least 3 separate experiments (N), and each experiment was done in triplicates (n=3) unless specified otherwise in text. Data were analyzed by 2-tail Student’s *t*-test or 1-way ANOVA, or 2-way ANOVA followed by Tukey’s post hoc multiple comparisons tests, where noted. Analysis was performed with assistance of GraphPad Prism (GraphPad, La Jolla, CA) software package. Significance was set at p < 0.05.

## Results

### Hippocampal delivery of AAV-NF-α1/CPE and AAV-NF-α1/CPE-E342Q rescues memory deficits in 3xTg-AD mice

3xTg-AD and non-Tg mice received bilateral hippocampal injections of either AAV-GFP, NF-α1/CPE, or NF-α1/CPE-E342Q at age 2 months (presymptomatic) and they then underwent a series of behavioral tests at age 9 months when cognitive decline was evident (Fig1A). Mice were first evaluated for locomotor activity in open field test. There were no significant differences across all four groups in travel distance and speed (Fig1B,1C). In the Morris Water-maze test, NF-α1/CPE and NF-α1/CPE-E342Q treated mice exhibited a trend of decrease in the latency of learning curve (Fig1D); and effectively increased the time spent in target quadrant on probe test in comparision to 3xTg+GFP mice (Figure1.E). All four groups of mice did not show any significant differences in swimming speed and distance on probe test of Morris water maze test (Fig1F,G).

**Figure 1.**
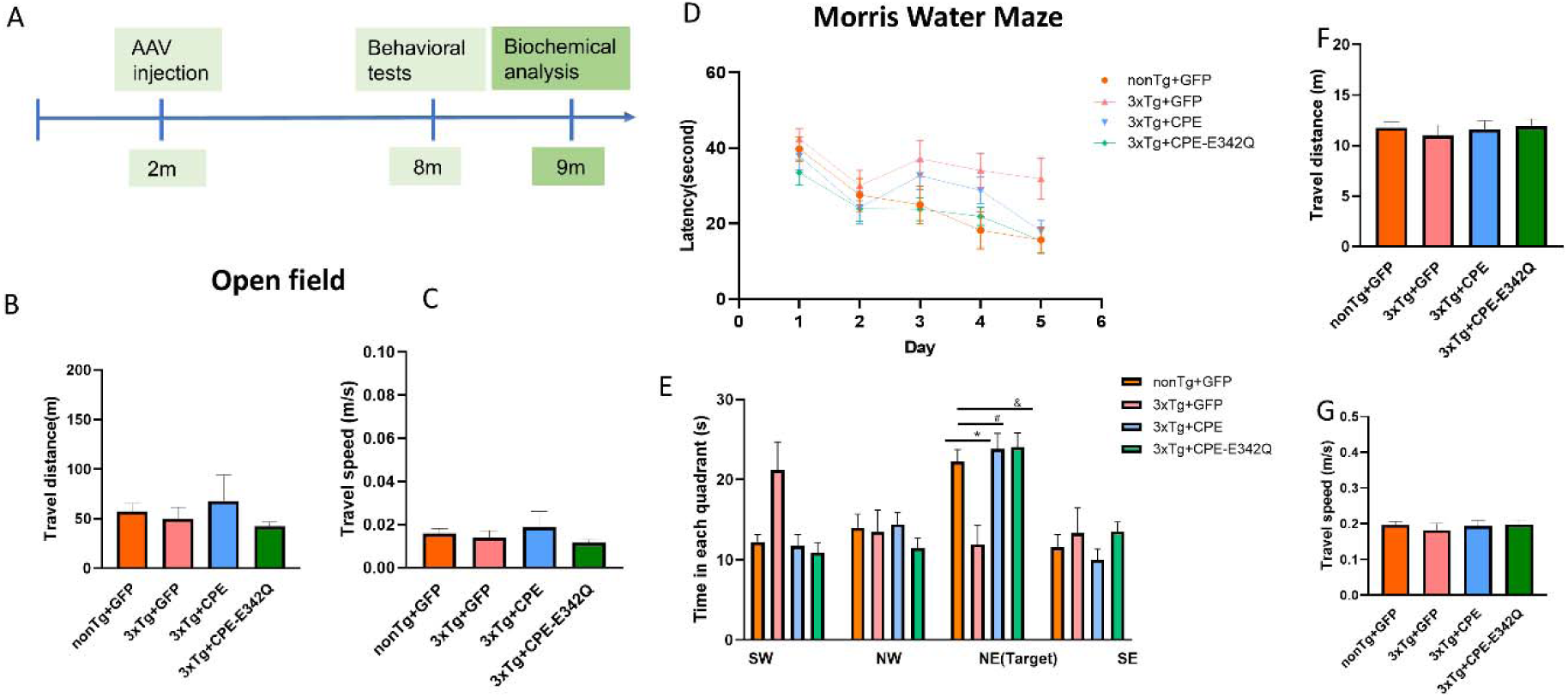
A. Schematic graph of experimental design. 3xTg-AD and nonTg mice received bilateral hippocampal injections of AAV-GFP or AAV-NF-α1/CPE or AAV-NF-α1/CPE-E342Q at age of 2.5 months and were evaluated by a series of behavioral tests at age of 8-9 months. Pathology of the hippocampus of the mice was examined at age of ∼9 months after behavioral tests. B, C. Locomotor activity in open field test. There were no significant differences across the groups in the travel distance and speed in the open field test. D, E. The effect of overexpression of NF-α1/CPE or NF-α1/CPE-E342Q on spatial learning in Morris water maze test. D. 3xTg+GFP mice displayed a trend of longer latency compared with other groups from day 1 to day 5. E. Overexpression of NF-α1/CPE or NF-α1/CPE-E342Q prevented memory deficit of 3xTg-AD mice in Morris water maze test. 3xTg+GFP mice spent less time in the target area (NE) in comparison with nonTg+GFP (*p=0.0232). While both 3xTg+ NF-α1/CPE (# p=0.0022) and 3xTg+ NF-α1/CPE-E342Q (& p=0.0015) mice spent more time in the target quadrant in comparison with 3xTg+GFP mice. One-way ANOVA analysis followed by Tukey’s post-hoc multiple comparison test [F(15,152) =6.566, p<0.0001]. n=10-11. Values are mean ± SEM. F, G Travel distance and speed in Morris water maze test. There were no significant differences in swimming distance or speed across all groups. n=10-11. Values are mean ± SEM.

### AAV-NF-α1/CPE and AAV-NF-α1/CPE-E342Q treatment rescues hippocampal CA1 neurodegeneration and decreases activated microglia in 3xTg-AD mice

Western blotting showed that NF-α1/CPE and NF-α1/CPE-E342Q expression was increased in 3xTg -AD mice treated with AAV-NF-α1/CPE and AAV-NF-α1/CPE-E342Q in comparison with those treated with GFP (Fig2A). Hippocampal CA1 neurodegeneration is significantly associated with AD [26]. MAP2 immunostaining in the CA1 region showed neurodegeneration (Fig2B) and decreased MAP2 intensity in 3xTg+GFP mice in comparison with nonTg+GFP mice, while significantly increased in 3xTg+CPE and partially increased in 3xTg+CPE-E343Q mice (Fig2C).

To evaluate whether AAV-NF-α1/CPE and AAV-NF-α1/CPE-E342Q treatment have any effects on neuroinflammation, GFAP and CD68 immunostaining were used to assess activated astrocytes and microglia. GFAP positive cells were similar across all four groups (Fig2D.E), while CD68 positive cells were increased in 3xTg+GFP group in comparison with non-Tg and reduced in 3xTg+CPE and 3xTg+CPE-E342Q mice in comparison with 3xTg+GFP (Fig 2F,G). This result indicates that NF-α1/CPE and NF-α1/CPE-E342Q might be involved in regulating neuroinflammatory response via modulating microglia, rather than astrocyte activity.

**Figure 2.**
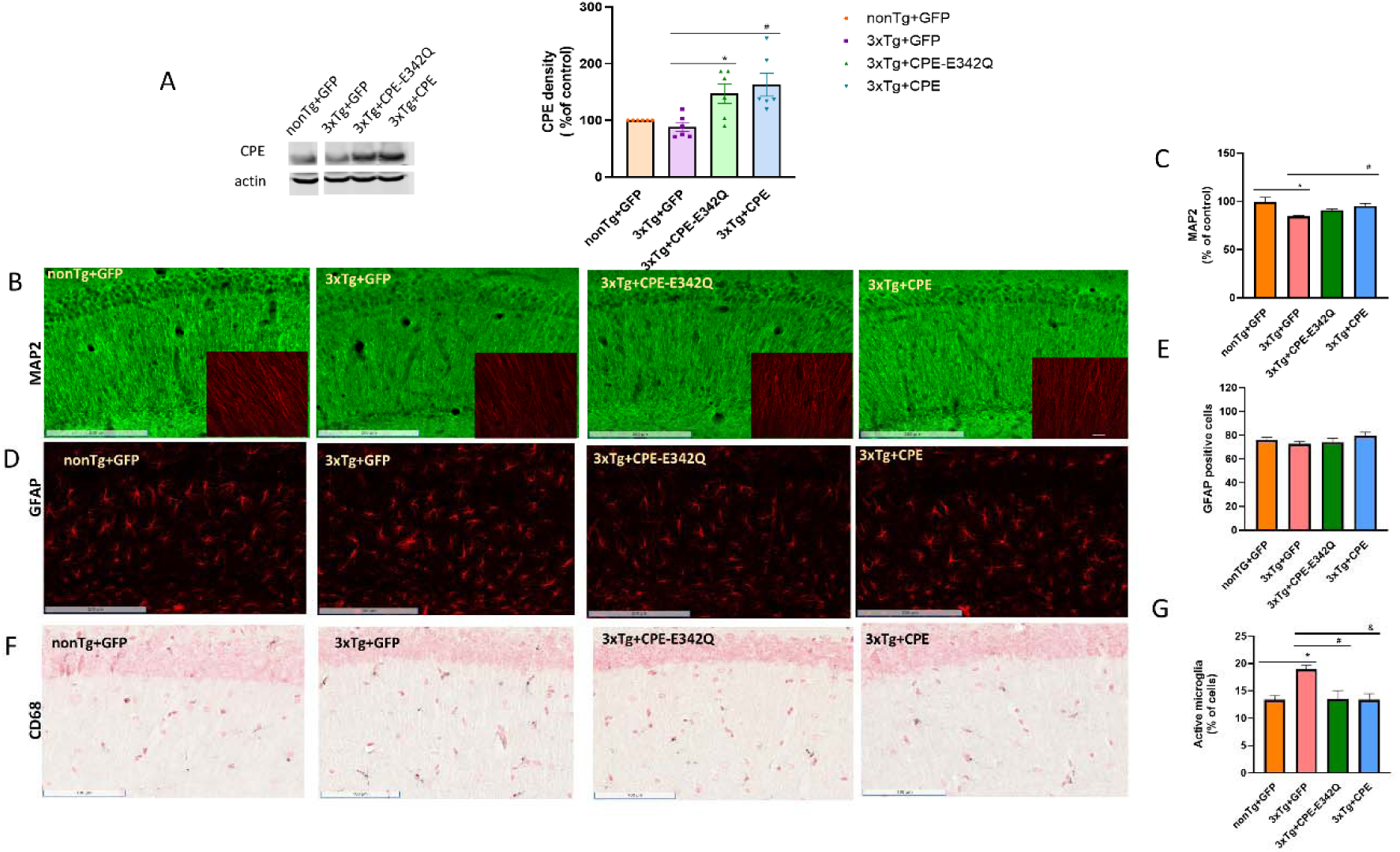
A. Effect of hippocampal stereotaxic injection of AAV-NF-α1/CPE and AAV-NF-α1/CPE-E342Q on CPE. CPE protein was increased in the hippocampus of 3xTg-AD mice that received AAV-NF-α1/CPE ( # p= 0.0059) and AAV-NF-α1/CPE-E342Q (* p=0.0351) treatment in comparison with 3xTg+GFP mice. One-way ANOVA analysis followed by Tukey’s post-hoc multiple comparison test [F(3,20) =6.643, p<0.01]. n=6, Values are mean ± SEM. B. Representative immunohistochemistry images of MAP2 of hippocampal CA1 region of nonTg+GFP, 3xTg+GFP, 3xTg+CPE-E342Q and 3xTg+CPE mice at age of 8-9 months. Scale bar =200 μm; Inset: scale bar =25 μm. n=6 mice per genotype. C. Quantification of MAP2 intensity in CA1 region of nonTg+GFP, 3xTg+GFP, 3xTg+CPE and 3xTg+CPE-E342Q mice at age of 8-9 months. MAP2 intensity decreased in 3xTg+GFP mice in comparison with nonTg+GFP, *p=0.0024; 3xTg+CPE increased MAP2 intensity significantly in comparison with 3xTg+GFP, # p=0.0284. One-way ANOVA analysis followed by Tukey’s post-hoc multiple comparison test, [F (3,17) = 6.871, p=0.0031]. Values are mean ± SEM. 3-4 sections per mouse, n=5-6 mice per genotype. nonTg+GFP made = 100% as control. D. Representative immunohistochemistry images of GFAP staining of hippocampal CA1 of nonTg+GFP, 3xTg+GFP, 3xTg+CPE-E342Q and 3xTg+CPE mice at age 8-9 months. Scale bar=200 μm. n=6 mice per genotype. E. Quantification of GFAP positive cells in CA1 region of nonTg+GFP, 3xTg+GFP, 3xTg+CPE-E342Q and 3xTg+CPE mice at age 8-9 months. There were no significant changes across all groups. One-way ANOVA analysis followed by Tukey’s post-hoc multiple comparison test. Values are mean ± SEM. 4 sections per mouse, n=6 mice per genotype. Values are mean ± SEM F. Representative immunohistochemistry images of CD68 immunostaining for activated microglia in hippocampal CA1 of nonTg+GFP, 3xTg+GFP, 3xTg+CPE and 3xTg+CPE-E342Q mice at age 8-9 months. Scale bar= 100 μm. Quantification of CD68 positive cells in hippocampal CA1 of nonTg+GFP, 3xTg+GFP, 3xTg+CPE-E342Q and 3xTg+CPE mice at age 8-9 months. CD68 positive cells were significantly increased in 3xTg+GFP in comparison with nonTg+GFP, *p=0.0141. Overexpression of NF-α1/CPE-E342Q and NF-α1/CPE in 3xTg-AD mice reduced activated microglia (CPE-E343Q: # p=0.0119; CPE:& p=0.0142). One-way ANOVA analysis followed by Tukey’s post-hoc multiple comparison test, [F(3,18) =6.196, p=0.0044]. 4 sections per mouse, n=5-6 mice per genotype. Values are mean ± SEM.

### AAV-NF-α1/CPE and AAV-NF-α1/CPE-E342Q decrease APP and tau phosphorylation and increase pro-survival protein, Bcl2, in 3xTg-AD mice

Similar to other AD mouse models, APP immunostaining showed strong intensity in the whole hippocampal area in 3xTg-AD mice, while it was very low in non-Tg mice (Fig3A). Some APP immunostaining was also observed in 3xTg+CPE and 3xTg+E342Q mice (Fig3A). Using an antibody specifically targeting human APP, Western blot showed that APP was decreased in 3xTg mice with both AAV-NF-α1/CPE-E342Q and AAV-NF-α1/CPE treatment (Fig3B). Studies using an antibody targeting both mouse and human APP showed that APP was much higher in 3xTg+GFP mice in comparison with nonTg+GFP mice, and treatment with AAV-NF-α1/CPE-E342Q and AAV-NF-α1/CPE effectively reduced APP levels in 3xTg mice (Fig3C). In addition, hippocampal AAV-NF-α1/CPE-E342Q or AAV-NF-α1/CPE delivery led to a decrease in the ratio of insoluble (Fig 3E), but not soluble amyloid 42/40 (Fig 3D).

**Figure 3.**
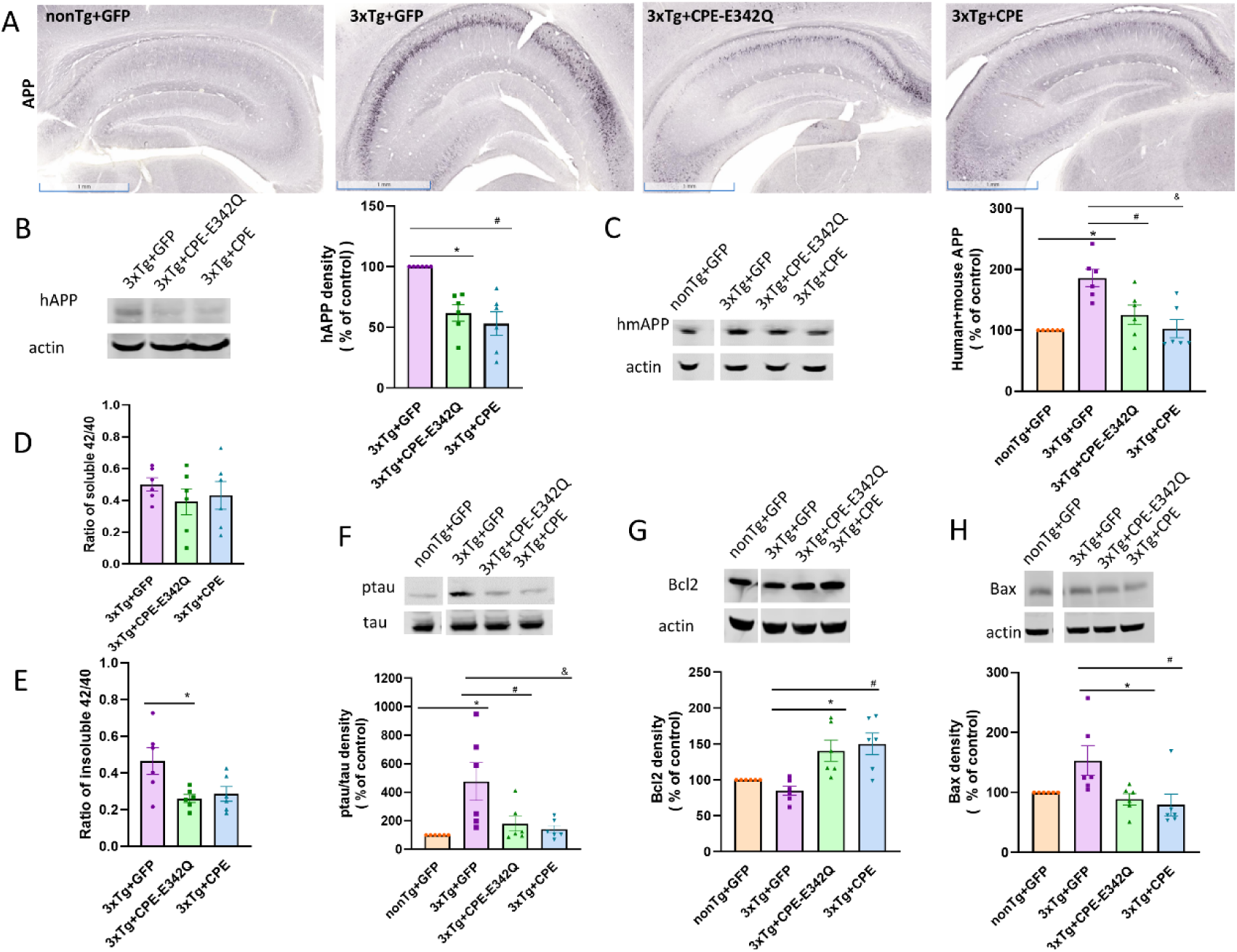
A. Representative immunohistochemistry images of APP expression after hippocampal stereotaxic injection in nonTg+GFP, 3xTg+GFP, 3xTg+CPE and 3xTg+CPE-E342Q mice at 8-9 months of age. Scale bar =1mm. B. Effect of hippocampal stereotaxic injection of AAV-NF-α1/CPE and AAV-NF-α1/CPE-E342Q on human APP(hAPP). hAPP was decreased by treatment with AAV-NF-α1/CPE and AAV-NF-α1/CPE-E342Q (CPE-E342Q: * p=0.0037; CPE: # p=0.0006;). One-way ANOVA analysis followed by Tukey’s post-hoc multiple comparison test [F(2,15) =13.16, p=0.0005]. n=6, Values are mean ± SEM. C. Effect of hippocampal stereotaxic injection of AAV-NF-α1/CPE and AAV-NF-α1/CPE-E342Q on human+mouse APP(hmAPP). hmAPP was significantly increased in 3xTg+GFP in comparison with nonTg+GFP ( * p=0.0008), while treatment with AAV-CPE and AAV-CPE-E342Q decreased hmAPP in 3xTg-AD mice (CPE-E342Q: # p=0.0186; CPE: & p=0.0012). One-way ANOVA analysis followed by Tukey’s post-hoc multiple comparison test [F(3,20) =9.301, p=0.0005]. n=6, Values are mean ± SEM. D. Effect of hippocampal stereotaxic injection of AAV-NF-α1/CPE and AAV-NF-α1/CPE-E342Q on ratio of soluble42/40. Treatment with AAV-CPE and AAV-E342Q did not induce any significant changes in ratio of soluble 42/40. n=6, Values are mean ± SEM. E. Effect of hippocampal stereotaxic injection of AAV-NF-α1/CPE and AAV-NF-α1/CPE-E342Q on the ratio of insoluble 42/40. Treatment with AAV-CPE-E342Q significantly decreased the ratio of insoluble 42/40 in 3xTg-AD, and AAV-CPE induce a trend of reduction in the ratio of insoluble 42/40 (CPE-E342Q: *p=0.0285; CPE: p=0.0555;). n=6, Values are mean ± SEM. F. Effect of hippocampal stereotaxic injection of AAV-NF-α1/CPE and AAV-NF-α1/CPE-E342Q on ratio of pTau/Tau. Ratio of pTau/Tau was significantly increased in 3xTg+GFP mice in comparison with nonTg+GFP ( * p=0.0078), while treatment with AAV-CPE and AAV-CPE-E342Q decreased ratio of pTau/Tau in 3xTg-AD (CPE-E342Q: # p=0.042;CPE: & p=0.0184,). One-way ANOVA analysis followed by Tukey’s post-hoc multiple comparison test [F(3,20) =5.555, p=0.0061]. n=6, Values are mean ± SEM. G. Effect of hippocampal stereotaxic injection of AAV-NF-α1/CPE and AAV-NF-α1/CPE-E342Q on Bcl2. Treatment with AAV-CPE and AAV-CPE-E342Q increased Bcl2 in 3xTg-AD mice (CPE-E342Q: * p=0.0084; CPE: # p=0.0021;). One-way ANOVA analysis followed by Tukey’s post-hoc multiple comparison test [F(3,20) =8.318, p=0.0009]. n=6, Values are mean ± SEM. H. Effect of hippocampal stereotaxic injection of AAV-NF-α1/CPE and AAV-NF-α1/CPE-E342Q on Bax. Treatment with AAV-CPE and AAV-CPE-E342Q decreased Bax in 3xTg-AD mice (CPE-E342Q: * p=0.0449;CPE: # p=0.0184;). One-way ANOVA analysis followed by Tukey’s post-hoc multiple comparison test [F(3,20) =4.274, p=0.0174]. n=6, Values are mean ± SEM.

Western blot analysis of tau phosphorylation revealed highly increased levels of ptau in 3xTg+GFP mice compared with nonTg+GFP mice. However, AAV-NF-α1/CPE-E342Q or AAV-NF-α1/CPE treatment significantly decreased ptau in these mice (Fig3 F).

Mitochondrial pro-survival protein, Bcl2 levels, is downregulated within tangle-bearing neurons of AD patients [27]; while expression of Bax, a mitochondrial pro-apoptotic protein is upregulated in AT8 positive cells, a marker for hyperphosphorylated Tau, in AD patients [28]. In 3xTg-AD mice, Bcl2 and Bax levels were not significantly different between nonTg+GFP and 3xTg+GFP, but treatment with AAV-NF-α1/CPE-E342Q or AAV-NF-α1/CPE increased Bcl2 (Fig3G) and decreased Bax (Fig3H), indicating the effect of NF-α1/CPE on enhancing the Bcl2-mediated pro-survival cascade.

### Proteomic analysis reveals differentially expressed proteins in hippocampus of AAV-NF-α1/CPE versus AAV-GFP treated 3xTg-AD mice

Proteome profiling analysis of hippocampus tissue from nonTg+GFP, 3xTg+GFP and 3xTg+ CPE mice was performed with LC-MS/MS as described in methods and illustrated in Fig4A. Volcano plots (Fig. 4B) show that there were 2814 differentially expressed proteins (DEPs) identified. The list of proteins with fold enrichment and p-values are provided in Supplementary Table S3. 3xTg+GFP in comparison with nonTg+GFP mice showed asymmetric distribution with majority of proteins showing higher expression (middle, magenta dots). In comparison, 3xTg+CPE mice compared to nonTg+GFP mice showed a more symmetric volcano plot (right, cyan dots). Notably, there were more DEPs down-regulated than up-regulated with AAV-NF-α1/CPE treatment of 3xTg-AD mice (left panel, grey dots). Comparable proteomic signatures were observed across replicates within three groups as shown by protein profile hierarchical heatmap (Fig. 4C). The heatmap data also shows the general increase in protein expression in 3xTg+GFP mice (red color) is reversed in 3xTg+CPE mice making it look comparable to nonTg+GFP mice (green color). These results suggest that AAV-NF-α1/CPE treatment reversed the global protein changes associated with AD pathology.

**Figure 4.**
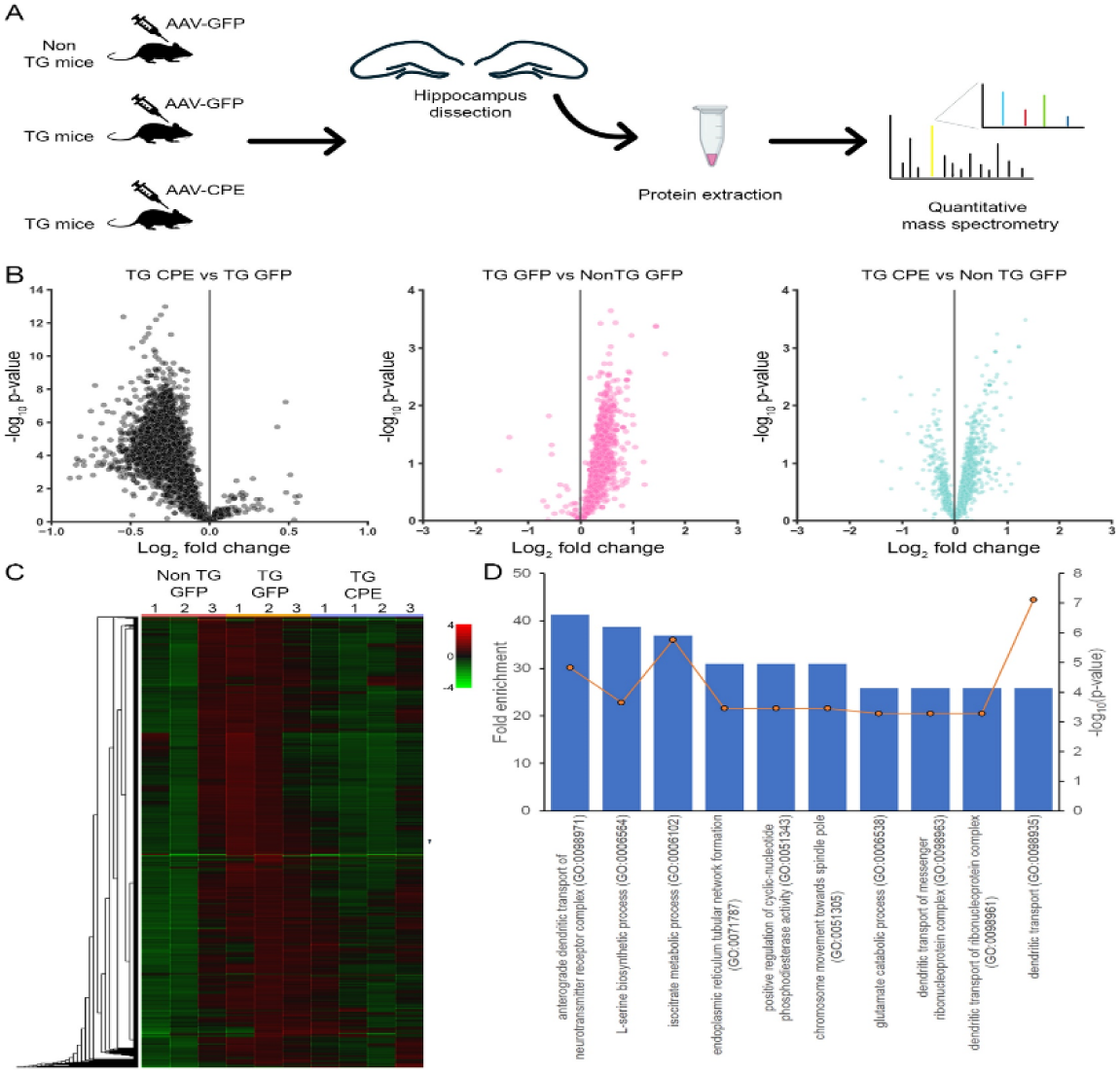
A. Quantitative mass spectrometry was performed on proteins extracted from hippocampus dissected from 3 groups of mice: nonTg+GFP, 3xTg+GFP, 3xTg+CPE in triplicates. B. Volcano plot with log_2_ fold change (x-axis) and -log_10_ p-value (y-axes) showing quantitative comparison of hippocampal proteins between 3xTg+CPE vs 3xTg+GFP, 3xTgGFP vs nonTg+GFP mice (magenta), 3xTg+ CPE vs nonTg+GFP mice (cyan). C. Protein profile hierarchical heatmap showing protein expression across the hippocampal samples from 3 comparison groups and their individual replicates (marked as 1, 2 and 3). The cell color represents log_2_ (ratio) to the average abundance across different samples. D. Top 10 canonical pathways from PANTHER Overrepresentation Test analysis using the dataset of most changed proteins filtered with log_2_ fold change ratio ≥ 0.33 or ≤ -0.33 between hippocampal proteins from 3xTg+ CPE vs 3xTg+GFP mice. Blue bars, associated with the left y axis, show fold enrichment, and circle orange markers, associated with the right y-axis, show the significance values plotted as −log (P-value).

Analysis of PANTHER Overrepresentation Test (Fisher’s Exact test) against the Gene Ontology molecular function dataset comparing 3xTg+CPE mice vs 3xTg+GFP mice showed enrichment in anterograde dendritic transport of neurotransmitter receptor complex, glutamate catabolic process, dendritic transport of messenger ribonucleoprotein complex, dendritic transport of ribonucleoprotein complex, and dendritic transport (Fig.4D), indicting AAV-NF-α1/CPE overexpression triggered synaptic and dendritic remodeling.

### Hippocampal delivery of AAV-NF-**α**1/CPE modulates expression of many AD-associated differentially expressed proteins in 3xTg-AD mice

STRING analysis of dataset comparing 3xTg+CPE mice vs 3xTg+GFP mice provided Alzheimer disease (KEGG ID: mmu05010) as one of the major KEGG network, with 83 proteins in dataset associated with Alzheimer’s disease network (Fig 5A). Further STRING network analysis of AD associated proteins in the dataset showed ones that are involved in mitochondrial ATP synthesis (Fig 5A, green), proteasomal protein catabolic process (Fig 5A, red), mitochondrial ATP synthesis electron transport (Fig 5A, purple), and regulation of autophagy (Fig 5A, magenta). Among the network of DEPs associated with AD regulated by NF-α1/CPE in 3xTg-AD mice, two new proteins, Trim28 and SNX4 were uncovered. Trim28 and Snx4 have been reported to have effects on promoting Tau and Aβ pathology in AD mice, respectively. Quantitative proteomic analysis showed that Trim28 and SNX4 were downregulated with expression of CPE in the hippocampus of 3xTg mice. To validate the findings of the proteomic data, we carried out Western blotting to analyze protein levels of Trim28 and SNX4 in the hippocampus of non-Tg and 3xTg-AD mice. Treatment with AAV-NF-α1/CPE or AAV-NF-α1/CPE-E342Q significantly decreased Trim28 levels in 3xTg+CPE and 3xTg+CPE-E342Q mice in comparison with 3xTg+GFP mice (Fig 5B). Similarly, SNX4 was decreased with AAV-NF-α1/CPE and AAV-NF-α1/CPE-E342Q treatment in 3xTg-AD mice compared to 3xTg+GFP mice (Fig5C).

**Figure 5.**
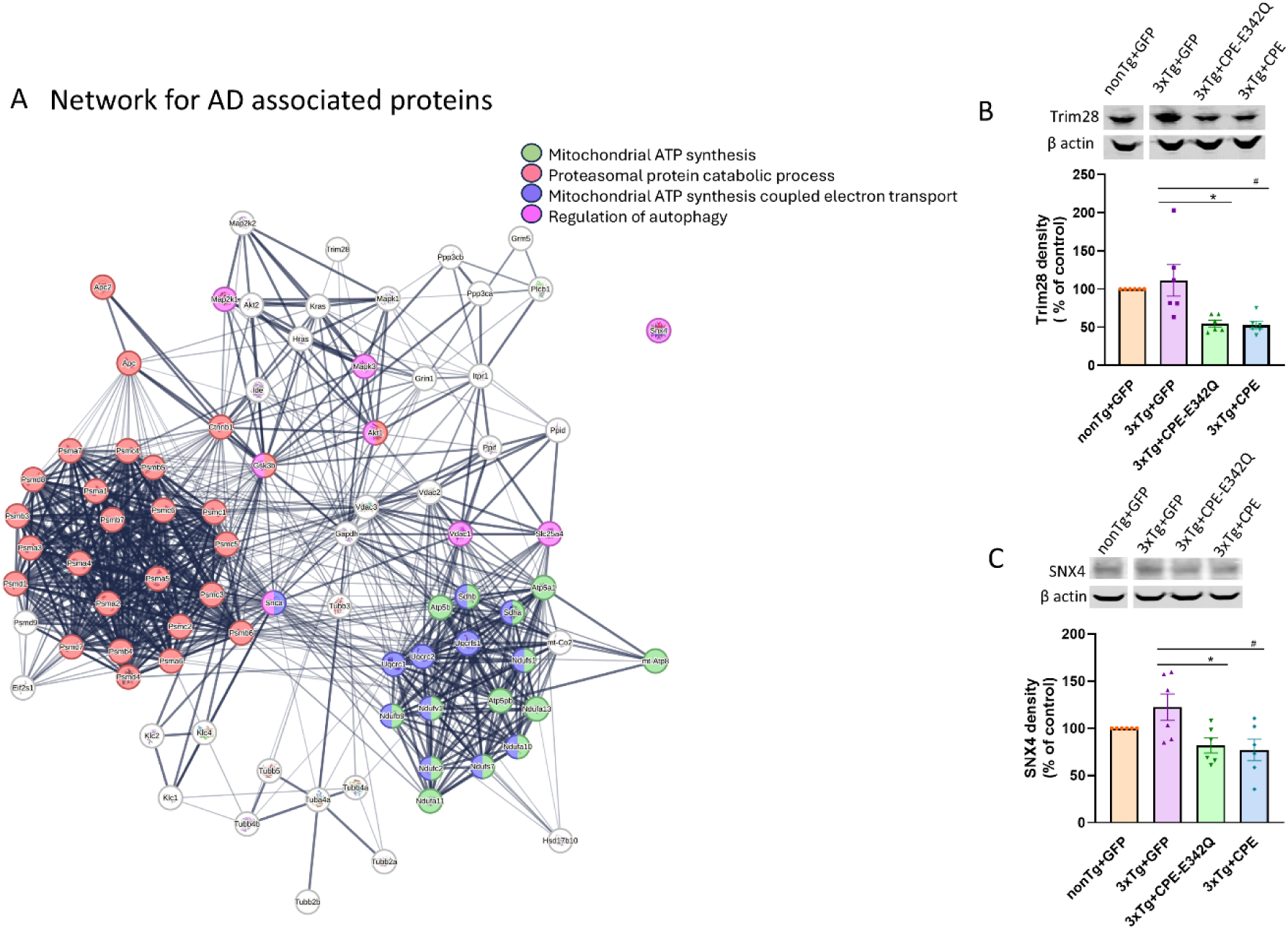
A. The functional protein network association maps were generated using data quantitatively comparing hippocampal proteins from 3xTg+ CPE vs 3xTg+GFP mice using String db. Signaling protein network of proteins annotated to be involved in Alzheimer’s disease (AD). The subset of proteins annotated to be involved in mitochondrial ATP synthesis (green), proteasomal protein catabolic process (red), mitochondrial ATP synthesis coupled protein transport (purple) and regulation of autophagy (magenta) are highlighted. B. Immunoblot and quantification of effects of AAV-NF-α1/CPE and AAV-NF-α1/CPE-E342Q on Trim28. Treatment with AAV-NFα1/CPE or AAV-CPE-E342Q decreased Trim28 in 3xTg-AD mice (CPE-E342Q: * p=0.0073; CPE: # p=0.0057). One-way ANOVA analysis followed by Tukey’s post-hoc multiple comparison test, [F (3,20) = 7.777, p=0.0012]. n=6 mice per genotype. Values are mean ± SEM. C. Immunoblot and quantification of effects of AAV-NF-α1/CPE and AAV-NF-α1/CPE-E342Q on SNX4. Treatment with AAV-NFα1/CPE or AAV-CPE-E342Q both decreased SNX4 in 3xTg-AD mice(CPE-E342Q: * p=0.0384; CPE: # p=0.0187). One-way ANOVA analysis followed by Tukey’s post-hoc multiple comparison test, [F (3,20) = 4.399, p=0.0157]. n=6 mice per genotype. Values are mean ± SEM.

### Hippocampal AAV-NF-**α**1/CPE delivery modulates synaptic proteins and rescues impaired synaptogenesis in 3xTg-AD mice

Synapse loss is highly correlated with cognitive impairments in AD patients [29–31]. In accord with findings in human, decreased synaptic density and activity were also observed in AD mouse models [32, 33]. STRING pathway analysis of dataset comparing expression between 3xTg+CPE mice vs 3xTg+GFP mice revealed that AAV-NF-α1/CPE treatment modulates expression of multiple proteins involved in synaptogenesis in 3xTg-AD mice (Fig 6A). Further analysis of synaptic organization network revealed the network that are primarily involved in postsynaptic density organization (Fig 6A, green), regulation of synaptic plasticity(Fig 6A, red), and postsynaptic organization (Fig 6A, purple).

**Figure 6.**
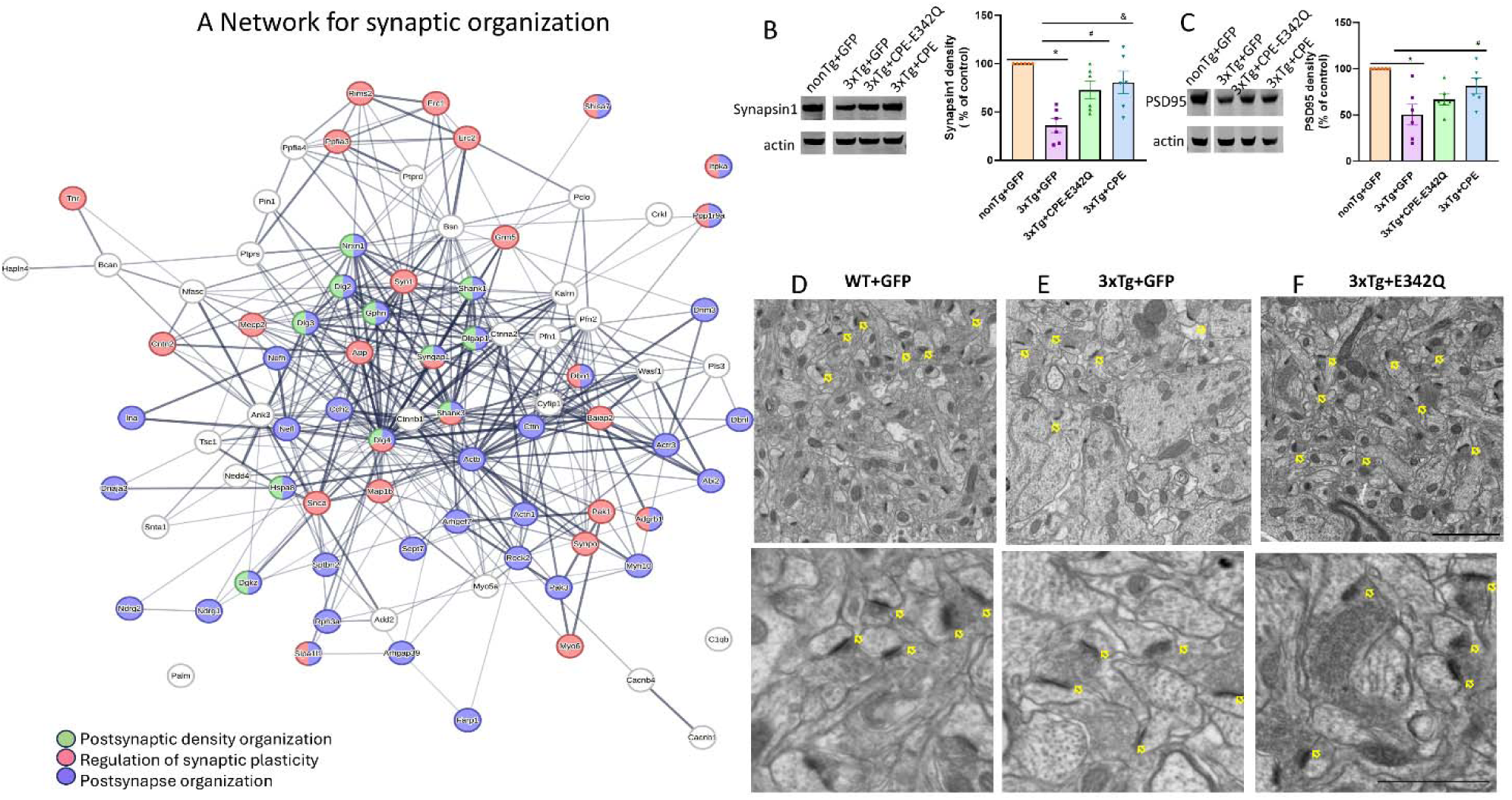
A. The functional protein network association maps were generated using data quantitatively comparing hippocampal proteins from 3xTg+ NF-α1/CPE vs 3xTg+GFP mice using String db. Signaling protein network of proteins annotated to be involved in synaptic organization. The subset of proteins annotated to be involved in postsynaptic density organization (green), regulation of synaptic plasticity (red) and postsynaptic organization (purple) are highlighted. B. Effects of AAV-NF-α1/CPE and AAV-NF-α1/CPE-E342Q on synaptic marker Synapsin1. Synapsin1 was significantly decreased in 3xTg+GFP mice in comparison with nonTg+GFP *p=0.0001. treatment with AAV-NFα1/CPE and AAV-CPE-E342Q effectively increased synpasin1 in 3xTg-AD mice ( CPE-E342Q:# p=0.0239; CPE:& p=0.0055. One-way ANOVA analysis followed by Tukey’s post-hoc multiple comparison test, [F (3,20) = 10.52, p=0.0002]. n=6 mice per genotype. Values are mean ± SEM. C. Effects of AAV-NF-α1/CPE and AAV-NF-α1/CPE-E342Q on synaptic marker PSD95. PSD95 was significantly decreased in 3xTg+GFP mice in comparison with nonTg+GFP *p=0.001. Treatment with AAV-NFα1/CPE increased PSD95 in 3xTg-AD mice, # p=0.0454. One-way ANOVA analysis followed by Tukey’s post-hoc multiple comparison test, [F (3,20) = 7.573, p=0.0014]. n=6 mice per genotype. Values are mean ± SEM. D.E.F. Electron microscopic image of nonTg+GFP (D), 3xTg+GFP (E) and 3xTg+CPE-E342Q (F). arrow: synapse. scale bar=2um upper panel; scale bar= 1um bottom panel.

To test the hypothesis provided by our proteomic analysis, we investigated two major synaptic proteins Synapsin1 and PSD95. Western blot analysis of Synapsin1 and PSD95 showed significant decrease in 3xTg+GFP mice in comparison with nonTg+GFP (Fig 6 B, C). Synapsin1 is mainly distributed in the vesicles of presynaptic terminals and regulates neurotransmitter release [34–36]. PSD95 is a critical component of post-synaptic density that interacts with many post synaptic proteins. Treatment with AAV-NF-α1/CPE significantly upregulated expression of Synapsin1 (Fig 6B) and PSD95(Fig 6C) in 3xTg+CPE mice. AAV-NF-α1/CPE-E342Q treatment increased Synapsin1 levels and induced a trend of increase of PSD95 in 3xTg+CPE-E342Q comparison with 3xTg+GFP mice (Fig 6C). Electron microscopic study showed synapses in the hippocampus of nonTg+GFP (Fig 6D) and 3xTg+CPE-E342Q (Fig 6F) mice have thick post synaptic thickening, in contrast to the thin post synaptic structures in the 3xTg+GFP mice (Fig 6E). This finding suggest defective synaptogenesis in 3xTg-AD mice is reversed with AAV-NF-α1/CPE or AAV-NF-α1/CPE-E342Q treatment.

### AAV-NF-**α**1/CPE and AAV-NF-α1/CPE-E342Q treatment restore autophagic activity in 3xTg-AD mice

Autophagy plays a crucial role in the pathogenesis of neurodegeneration [37]. Autophagosomes containing large amounts of dense core materials accumulated in the dystrophic neurites around amyloid in the cortex have been reported in post-mortem brain of patients with AD, linking impaired autophagy with AD [38]. Our proteomic analysis revealed a network of differentially expressed proteins with **AAV**-NF-α1/CPE treatment that are associated with autophagy pathways (Fig 7A). STRING analysis of dataset comparing 3xTg+CPE mice vs 3xTg+GFP mice revealed 135 autophagy associated proteins were altered.

**Figure 7.**
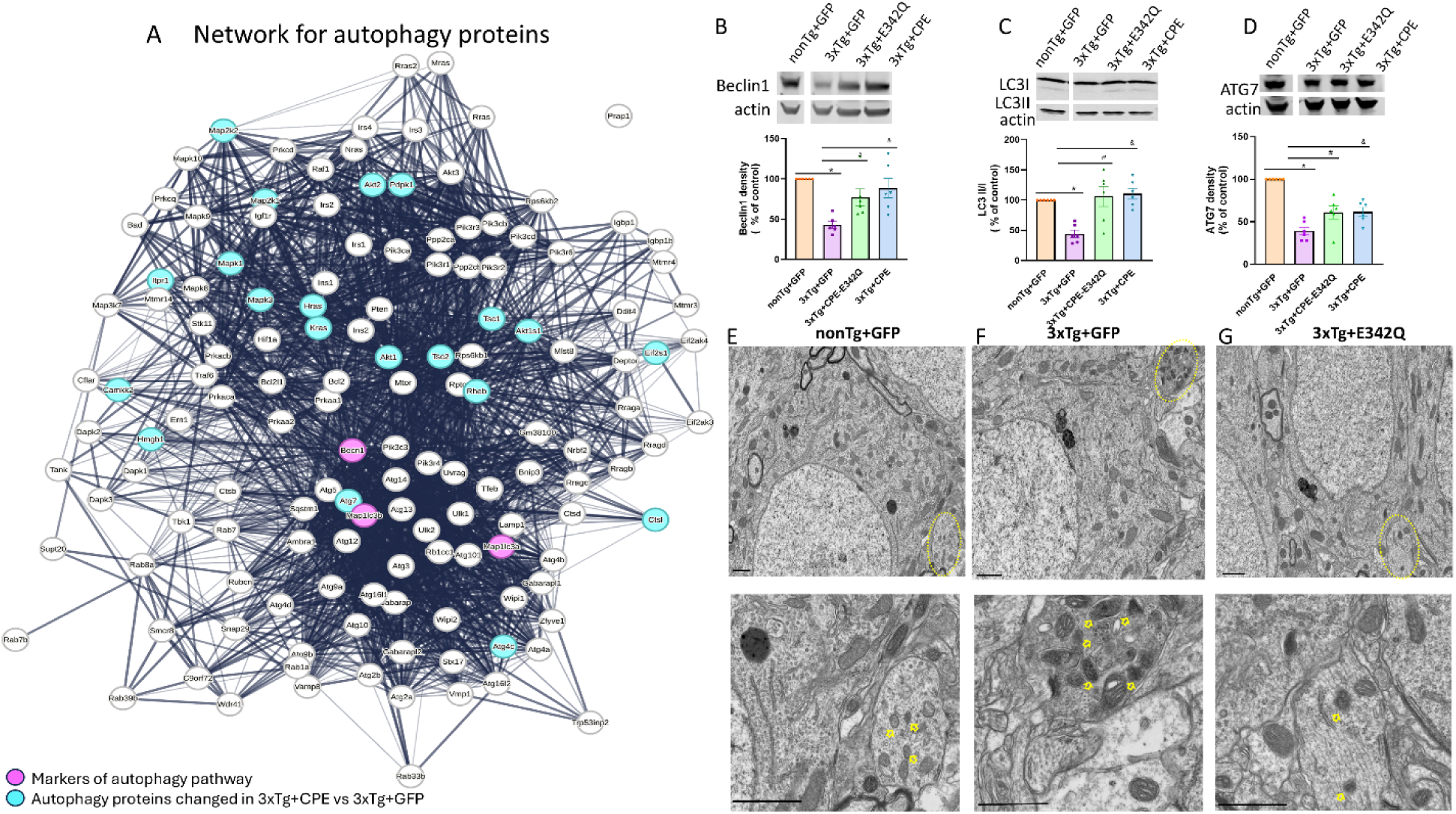
A. The String db functional protein network association map of autophagy related proteins. The proteins highlighted in cyan were detected in our experimental dataset comparing hippocampal proteins from 3xTg+CPE vs 3xTg+GFP mice. Beclin 1 and Map1lc3a/b (LC3 II/I) labeled in magenta are prominent markers of autophagy pathway. B. Immunoblot and quantification of effects of AAV-NF-α1/CPE and AAV-NF-α1/CPE-E342Q on Beclin1. Beclin1 was significantly decreased in 3xTg+GFP mice in comparison with nonTg+GFP *p=0.0005. Treatment with AAV-NFα1/CPE and CPE-E342Q both increased Beclin1 in 3xTg-AD mice (CPE-E342Q: # p=0.0426; CPE: & p=0.0049). One-way ANOVA analysis followed by Tukey’s post-hoc multiple comparison test, [F (3,20) = 8.743, p=0.0007]. n=6 mice per genotype. Values are mean ± SEM. C. Immunoblot and quantification of effects of AAV- NF-α1/CPE and AAV-NF-α1/CPE-E342Q on LC3II. Ratio of LC3II/LC3I was significantly decreased in 3xTg+GFP mice in comparison with nonTg+GFP *p=0.0041. Treatment with AAV-NFα1/CPE and CPE-E342Q both increased ratio of LC3II/LC3I in 3xTg-AD mice (CPE-E342Q: # p=0.0015; CPE: &p=0.0008) and. One-way ANOVA analysis followed by Tukey’s post-hoc multiple comparison test, [F (3,20) = 9.579, p=0.0004]. n=6 mice per genotype. Values are mean ± SEM. D. Effects of AAV-NF-α1/CPE and AAV-NF-α1/CPE-E342Q on autophagy-related protein ATG7. ATG7 was significantly decreased in 3xTg+GFP mice in comparison with nonTg+GFP *p<0.0001. Treatment with AAV-NFα1/CPE and CPE-E342Q both increased ATG7 in 3xTg-AD mice ( CPE-E342Q: # p=0.0347; CPE: & p=0.0309) One-way ANOVA analysis followed by Tukey’s post-hoc multiple comparison test, [F (3,20) =23.71, p<0.0001]. n=6 mice per genotype. Values are mean ± SEM. E. F.G. Electron microscopic image of nonTg+GFP (C), 3xTg+GFP (D) and 3xTg+E342Q (E). Circle: neurite (defined by the structure of synapsis) containing autophagosomes. Arrowhead: autophagosomes containing dense core material. Scale bar =1um.

To examine autophagy-related proteins that may be modulated by CPE or CPE-E342Q to rescue impaired autophagy in 3xTg-AD mice, immunoblot and electron microscopy (EM) studies were carried out. Beclin 1 and Map1lc3a/b (LC3 II/I) are established autophagy markers. Beclin1 is a protein that facilitates the assembly of autophagosomes via interacting with other proteins such as phosphoinositide 3-kinase (PI3KC3)/Vps34 [39, 40]. Western blots showed that Beclin1 was significantly decreased in 3xTg+GFP group in comparison with nonTG+GFP; while treatment with AAV-NF-α1/CPE and AAV-NF-α1/CPE-E342Q increased Beclin1 compared to the 3xTg+GFP group (Fig 7B). LC3II is a modified form of microtubule-associated light chain 3(LC3I) protein which is recruited to autophagosome membrane and later degraded by lysosomal hydrolases [41]. Western blot determination of the ratio of LC3II/LC3I showed that it was decreased in 3xTg-AD compared to non-Tg mice, and increased in 3xTg-AD mice with AAV-NF-α1/CPE and AAV-NF-α1/CPE-E342Q treatment (Fig.7C). The network analysis revealed the autophagy pathway protein, ATG7 downstream of Map1lc3a/b (LC3 II/I) and hence we examined its regulation of expression by NF-α1/CPE. We demonstrated by Western blot that the ATG7 level was reduced in 3xTg-AD mice, but enhanced with AAV-NF-α1/CPE and AAV-NF-α1/CPE-E342Q (Fig.7D).

EM studies show that autophagosomes in normal neurites in the hippocampus of nonTg+GFP mouse have little dense material (Fig 7E); in contrast, autophagosomes in 3xTg+GFP mouse contain highly dense undigested materials observed in dystrophic neurites (Fig 7F). In 3xTg+ CPE-E342Q mice, autophagosomes in neurites contained less dense material accumulated compared to 3xTg+GFP (Fig 7G). These results indicate that AAV-NF-α1/CPE and AAV-NF-α1/CPE-E342Q treatment rescues impaired autophagy in 3xTg-AD mice by up-regulating expression of critical proteins Beclin1, LC3 and ATG7.

## Discussion

Recent work from several laboratories have demonstrated that delivery of AAV or lentivirus - NF-α1/CPE or upregulation of NF-α1/CPE expression by agomirs can rescue AD pathology and cognitive decline in three different AD mouse models [13, 42, 43]. In the study by Fan et al [42], conditional knockout of NF-α1/CPE in 5xFAD mice across different ages resulted in enhancement of AD pathology. Furthermore, a direct dose-dependent correlation between CPE overexpression and decrease in amyloid plaque formation in the hippocampus was found [42]. In the study of Jiang et al, two agomirs administered through nasal instillation or via cerebral ventricular injection increased CPE expression which subsequenly enhanced neurogenesis in the dentate gyrus to reverse AD pathology and cognitive dysfunction. The proposed mechanism was based on their finding of increased CPE after agomir treatment which enhanced BDNF and FGF2 levels known to promote neurogenesis [43]. Our previous study has shown that hippocampal delivery of AAV-NF-α1/CPE into 3xTg-AD mice increased Bcl2 and decreased Bax, a mechanism known to mediate neuroprotection [13]. Additionally,Serpina3g, a pro-survival protein in mice was found to be up-regulated by AAV-NF-α1/CPE treatment [13]. Moreover, the AAV-NF-α1/CPE treated 3xTg-AD mice supressed the expression of APP through decreasing the transcription factors, Sp1 and Hsf1 which bind to the promoter of APP to regulate expression. A pro-inflammatory protein Card14 and a mitophagy inhibitor, Plin4, were downregulated in expression by the AAV-NF-α1/CPE treatment of 3xTg-AD mice to mitigate neuroinflammation and promote mitophagy, respectively. These previous studies have propelled NF-α1/CPE as a potentially important therapeutic agent for treatment of AD (Fig8).

In our current work, we have carried out a proteomic study which has uncovered novel proteins involved in AD pathogenesis and further illuminated the mechanisms underlying the actions of NF-α1/CPE in rescuing AD pathology and cognitive dysfunction in 3xTg-AD mice. These include restoring levels of proteins Synapsin1 and PSD95 that are involved in synaptogenesis which were decreased in 3xTg-AD mice (Fig 8). Analysis of DEPs in the autophagy pathway network followed by Western blotting revealed that NF-α1/CPE restored levels of Beclin1 and Map1lc3a/b (LC3II/1), and up-regulated the expression of its downstream protein ATG7. ATG7 is a crucial protein in the autophagy pathway, acting as an E1-like enzyme that initiates the process of autophagy and plays a vital role in forming and extending autophagosomes that engulf cellular waste and damaged components for recycling [44] (Fig 8). Furthermore, two proteins: Tripartite motif-containing 28 (Trim28) and SNX4 which showed aberrant levels in AD mice were decreased with NF-α1/CPE treatment. Trim 28 plays a critical role in regulating α-Syn and Tau levels. Reduction of Trim 28 decreased Tau- and α-Syn in drosophila [45] and mice [46]. Sorting nexin-4 (SNX4) regulates Aβ production. Reduction of SNX4 activity decreased the steady-state level of BACE1 and subsequent attenuation of Aβ production [47]. Thus, NF-α1/CPE acts in a multifaceted manner to restore cognitive dysfuntion and pathology present in AD mice.

**Figure 8.**
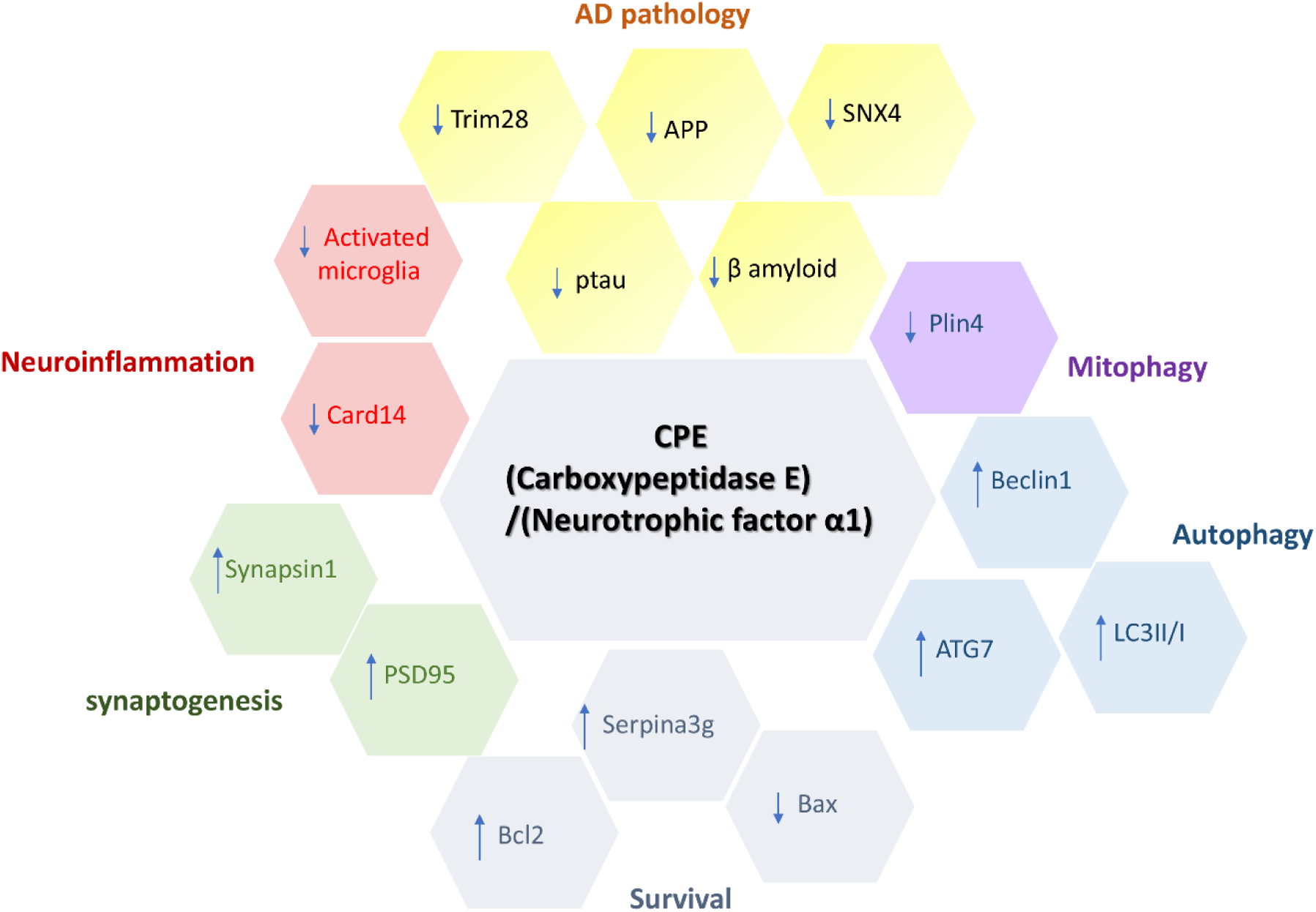
Summary of verified proteins regulated by NFα1/CPE in 3xTg-AD mice. NFα1/CPE plays a critical role in the pathogenesis of AD. It downregulates characteristic pathological proteins including APP, β amyloid and Ptau. Additionally, Trim28 and SNX4 which are involved in the production of pathological proteins are reduced by NFα1/CPE treatment. AD associated neuroinflammation, especially increased activation of microglia was reduced by NFα1/CPE treatment as evidence by decreased activated microglia number and Card14 protein. Autophagy markers, Beclin1, ATG7 and ratio of LC3II/I are all increased by NFα1/CPE treatment, indicating higher activity of autophagy. Synaptogenesis markers such as Synapsin1 and PSD95 are increased by NFα1/CPE, suggesting increased synaptogenesis. NFα1/CPE also acts as a neuroprotector by upregulating survival proteins. For instance, mitochondria survival protein, Bcl2 is enhanced, and pro-apoptotic protein, Bax is reduced by NFα1. Serpina3g, a protein that upregulates survival, is also increased by NFα1/CPE. Plin4, inhibitor of mitophagy, is reduced by NFα1/CPE. NFα1/CPE : Neurotrophic Factor-α1/carboxypeptidase E. APP: amyloid precursor protein. SNX4: sorting nexin 4. Trim28: tripartite motif containing 28. Card14; aspase recruitment domain-containing protein 14. PSD95: postsynaptic density protein 95. Serpina3g: Serine protease inhibitor A3G. Plin4: Perilipin 4. ATG7: autophagy related 7. LC3: Microtubule-associated protein 1A/1B-light chain 3.

Numerous proteomic studies using animal models and in AD patients have indicated that AD is a very complex disease, exhibiting dysregulation of multiple regulatory metabolic pathways [48–50]. An analysis of proteomic studies involving thirty-eight reports in clinical patients with AD revealed that synaptic-activity and vesicle associated proteins are altered in early stage of AD, and mitochondria function associated proteins are changed in more advanced stage of AD [51]. Aberrant autophagy has also been reported in AD patients [52]. Indeed, findings from our proteomic study in 3xTg mice mirror those from clinical studies. Dysregulation of vast numbers of proteins associated with mitochondrial activity, proteasomal protein catabolic process, dendritic transport and glutamate catabolic process and autophagy were also identified in our AD mice. Although there were no exact common dysregulated proteins identified in our 3xTg proteomic study with those from clinical AD proteomic analysis published thus far, there were many dysregulated proteins with common functions. For example, glutamate ionotropic receptor AMPA type subunit 2 (Gria2) was found decreased in AD patients [51], and Gria1 was also altered in the 3xTg mice with NF-α1/CPE treatment, (Fig S1) suggesting the role of AMPA subunit in the pathogenesis of AD. Changes of HspB1 in AD brains have been reported [51, 53] and alteration of Hspa1a was observed in 3xTg mice with NF-α1/CPE treatment (Fig S1). These findings indicate the involvement of Hsp families in the progression of AD. Hence, our proteomic study has uncovered new insights into expression of proteins that are altered in AD pathology and progression, providing an avenue for further translational and clinical studies, such as developing biomarkers for early detection of AD and following treatment efficacy.

## Conclusions

It is now highly recognized that AD is a multifactorial disease and yet, at present most therapeutic approaches have focused on single target such as elimination of Tau and amyloid accumulation in the brain, with only a small amount of success for long term reversal of cognitive dysfunction [54–57]. Moreover, these drugs give rise to many side effects [58]. Our current study together with previous work have strengthened the evidence that NF-α1/CPE is an excellent candidate therapeutic agent for treating AD and potentially other neurodegenerative diseases. Foremost we have demonstrated that NF-α1/CPE is different from other therapeutics in its ability to normalize and modulate protein expression in numerous cell biological processes, metabolic pathways, synaptogenesis and autophagy in neurons and glial cells which functions have been disrupted in AD, to regain homeostasis. Secondly, our proteomic study has uncovered new proteins linked to AD defects and insights into the mechanism of action of NF-α1/CPE in rescuing AD pathology. Finally, we have previously demonstrated that overexpression of NF-α1/CPE via AAV-gene therapy in normal mice has no apparent side effects on perturbing proteins disrupted in AD or changes in cognitive function [13].

## Acknowledgements

The authors thank Dr. Yogita Chudasama at Rodent Behavioral Core, NIMH, for their excellent training and advice on stereotaxic injection procedures and animal care after surgery and Chip Dye, NICHD Imaging Core for carrying out the electron microscopy.

## Author contributions

Y. P. Loh designed the research project; L. Xiao, X. Yang and D. Abebe performed the experimental research, P. Sharma performed the proteomic and bioinformatic study. Y. P. Loh, L. Xiao and P. Sharma analyzed the data; Y. P. Loh, L. Xiao and P. Sharma wrote the paper with contributions of various sections and editing from. X. Yang.

## Data availability

All data sets generated during and/or analyzed during the current study are available from the corresponding author on reasonable request.

## Declarations

### Ethics approval and consent to participate

All animal study protocols were approved by the Animal Care and Use Committee of NICHD, NIH and complied with ethical regulations.

### Consent for publication

All the authors listed in the article give their consent for the publication of this article.

## Abbreviations

AD: Alzheimer disease
AMPA: α-amino-3-hydroxy-5-methyl-4-isoxazolepropionic acid.
APP: Amyloid precursor protein.
ATG7: Autophagy related 7
Bdnf: Brain-derived neurotrophic factor.
Card14: Aspase recruitment domain-containing protein 14
Fgf2: Fibroblast growth factor 2
Gria: glutamate ionotropic receptor AMPA type subunits
Hsp: Heat shock protein
LC3: Microtubule-associated protein 1A/1B-light chain 3
NFα1/CPE: Neurotrophic Factor-α1/carboxypeptidase E
PI3KC3: Phosphoinositide 3-kinase
Plin4: Perilipin 4
PSD95: Postsynaptic density protein 95
Serpina3g: Serine protease inhibitor A3G
SNX4: sorting nexin 4
Trim28: Tripartite motif containing 28

## References

1. Collaborators GBDDF: Estimation of the global prevalence of dementia in 2019 and forecasted prevalence in 2050: an analysis for the Global Burden of Disease Study 2019. Lancet Public Health 2022, 7:e105–e125.

2. Mendez MF: Early-onset Alzheimer Disease and Its Variants. Continuum (Minneap Minn) 2019, 25:34–51.

3. Ballard C, Gauthier S, Corbett A, Brayne C, Aarsland D, Jones E: Alzheimer’s disease. Lancet 2011, 377:1019–1031.

4. Uddin MS, Stachowiak A, Mamun AA, Tzvetkov NT, Takeda S, Atanasov AG, Bergantin LB, Abdel-Daim MM, Stankiewicz AM: Autophagy and Alzheimer’s Disease: From Molecular Mechanisms to Therapeutic Implications. Front Aging Neurosci 2018, 10:04.

5. Yu WH, Cuervo AM, Kumar A, Peterhoff CM, Schmidt SD, Lee JH, Mohan PS, Mercken M, Farmery MR, Tjernberg LO, et al: Macroautophagy--a novel Beta-amyloid peptide-generating pathway activated in Alzheimer’s disease. J Cell Biol 2005, 171:87–98.

6. Dickstein DL, Brautigam H, Stockton SD, Jr., Schmeidler J, Hof PR: Changes in dendritic complexity and spine morphology in transgenic mice expressing human wild-type tau. Brain Struct Funct 2010, 214:161–179.

7. Polydoro M, Acker CM, Duff K, Castillo PE, Davies P: Age-dependent impairment of cognitive and synaptic function in the htau mouse model of tau pathology. J Neurosci 2009, 29:10741–10749.

8. Kopeikina KJ, Polydoro M, Tai HC, Yaeger E, Carlson GA, Pitstick R, Hyman BT, Spires-Jones TL: Synaptic alterations in the rTg4510 mouse model of tauopathy. J Comp Neurol 2013, 521:1334–1353.

9. About a peculiar disease of the cerebral cortex. By Alois Alzheimer, 1907 (Translated by L. Jarvik and H. Greenson). Alzheimer Dis Assoc Disord 1987, 1:3–8.

10. Hippius H, Neundorfer G: The discovery of Alzheimer’s disease. Dialogues Clin Neurosci 2003, 5:101–108.

11. Cheng Y, Cawley NX, Loh YP: Carboxypeptidase E/NFalpha1: a new neurotrophic factor against oxidative stress-induced apoptotic cell death mediated by ERK and PI3-K/AKT pathways. PLoS One 2013, 8:e71578.

12. Xiao L, Sharma VK, Toulabi L, Yang X, Lee C, Abebe D, Peltekian A, Arnaoutova I, Lou H, Loh YP: Neurotrophic factor-alpha1, a novel tropin is critical for the prevention of stress-induced hippocampal CA3 cell death and cognitive dysfunction in mice: comparison to BDNF. Transl Psychiatry 2021, 11:24.

13. Xiao L, Yang X, Sharma VK, Abebe D, Loh YP: Hippocampal delivery of neurotrophic factor-alpha1/carboxypeptidase E gene prevents neurodegeneration, amyloidosis, memory loss in Alzheimer’s Disease male mice. Mol Psychiatry 2023, 28:3332–3342.

14. Woronowicz A, Koshimizu H, Chang SY, Cawley NX, Hill JM, Rodriguiz RM, Abebe D, Dorfman C, Senatorov V, Zhou A, et al: Absence of carboxypeptidase E leads to adult hippocampal neuronal degeneration and memory deficits. Hippocampus 2008, 18:1051–1063.

15. Fan FC, Du Y, Zheng WH, Loh YP, Cheng Y: Carboxypeptidase E conditional knockout mice exhibit learning and memory deficits and neurodegeneration. Transl Psychiatry 2023, 13:135.

16. Durmaz A, Aykut A, Atik T, Ozen S, Ayyildiz Emecen D, Ata A, Isik E, Goksen D, Cogulu O, Ozkinay F: A New Cause of Obesity Syndrome Associated with a Mutation in the Carboxypeptidase Gene Detected in Three Siblings with Obesity, Intellectual Disability and Hypogonadotropic Hypogonadism. J Clin Res Pediatr Endocrinol 2021, 13:52–60.

17. Alsters SI, Goldstone AP, Buxton JL, Zekavati A, Sosinsky A, Yiorkas AM, Holder S, Klaber RE, Bridges N, van Haelst MM, et al: Truncating Homozygous Mutation of Carboxypeptidase E (CPE) in a Morbidly Obese Female with Type 2 Diabetes Mellitus, Intellectual Disability and Hypogonadotrophic Hypogonadism. PLoS One 2015, 10:e0131417.

18. Pla V, Paco S, Ghezali G, Ciria V, Pozas E, Ferrer I, Aguado F: Secretory sorting receptors carboxypeptidase E and secretogranin III in amyloid beta-associated neural degeneration in Alzheimer’s disease. Brain Pathol 2013, 23:274–284.

19. Sharma VK, Yang X, Kim SK, Mafi A, Saiz-Sanchez D, Villanueva-Anguita P, Xiao L, Inoue A, Goddard WA, 3rd, Loh YP: Novel interaction between neurotrophic factor-alpha1/carboxypeptidase E and serotonin receptor, 5-HTR1E, protects human neurons against oxidative/neuroexcitotoxic stress via beta-arrestin/ERK signaling. Cell Mol Life Sci 2021, 79:24.

20. Romberg C, Mattson MP, Mughal MR, Bussey TJ, Saksida LM: Impaired attention in the 3xTgAD mouse model of Alzheimer’s disease: rescue by donepezil (Aricept). J Neurosci 2011, 31:3500–3507.

21. Hutton M, Lendon CL, Rizzu P, Baker M, Froelich S, Houlden H, Pickering-Brown S, Chakraverty S, Isaacs A, Grover A, et al: Association of missense and 5’-splice-site mutations in tau with the inherited dementia FTDP-17. Nature 1998, 393:702–705.

22. Oddo S, Caccamo A, Shepherd JD, Murphy MP, Golde TE, Kayed R, Metherate R, Mattson MP, Akbari Y, LaFerla FM: Triple-transgenic model of Alzheimer’s disease with plaques and tangles: intracellular Abeta and synaptic dysfunction. Neuron 2003, 39:409–421.

23. Guo Q, Fu W, Sopher BL, Miller MW, Ware CB, Martin GM, Mattson MP: Increased vulnerability of hippocampal neurons to excitotoxic necrosis in presenilin-1 mutant knock-in mice. Nat Med 1999, 5:101–106.

24. Cheng Y, Cawley NX, Yanik T, Murthy SR, Liu C, Kasikci F, Abebe D, Loh YP: A human carboxypeptidase E/NF-alpha1 gene mutation in an Alzheimer’s disease patient leads to dementia and depression in mice. Transl Psychiatry 2016, 6:e973.

25. Belfiore R, Rodin A, Ferreira E, Velazquez R, Branca C, Caccamo A, Oddo S: Temporal and regional progression of Alzheimer’s disease-like pathology in 3xTg-AD mice. Aging Cell 2019, 18:e12873.

26. Bobinski M, de Leon MJ, Tarnawski M, Wegiel J, Reisberg B, Miller DC, Wisniewski HM: Neuronal and volume loss in CA1 of the hippocampal formation uniquely predicts duration and severity of Alzheimer disease. Brain Res 1998, 805:267–269.

27. Su JH, Satou T, Anderson AJ, Cotman CW: Up-regulation of Bcl-2 is associated with neuronal DNA damage in Alzheimer’s disease. Neuroreport 1996, 7:437–440.

28. Su JH, Deng G, Cotman CW: Bax protein expression is increased in Alzheimer’s brain: correlations with DNA damage, Bcl-2 expression, and brain pathology. J Neuropathol Exp Neurol 1997, 56:86–93.

29. Terry RD, Masliah E, Salmon DP, Butters N, DeTeresa R, Hill R, Hansen LA, Katzman R: Physical basis of cognitive alterations in Alzheimer’s disease: synapse loss is the major correlate of cognitive impairment. Ann Neurol 1991, 30:572–580.

30. Scheff SW, Price DA, Schmitt FA, DeKosky ST, Mufson EJ: Synaptic alterations in CA1 in mild Alzheimer disease and mild cognitive impairment. Neurology 2007, 68:1501–1508.

31. de Wilde MC, Overk CR, Sijben JW, Masliah E: Meta-analysis of synaptic pathology in Alzheimer’s disease reveals selective molecular vesicular machinery vulnerability. Alzheimers Dement 2016, 12:633–644.

32. Kashyap G, Bapat D, Das D, Gowaikar R, Amritkar RE, Rangarajan G, Ravindranath V, Ambika G: Synapse loss and progress of Alzheimer’s disease -A network model. Sci Rep 2019, 9:6555.

33. Hong S, Beja-Glasser VF, Nfonoyim BM, Frouin A, Li S, Ramakrishnan S, Merry KM, Shi Q, Rosenthal A, Barres BA, et al: Complement and microglia mediate early synapse loss in Alzheimer mouse models. Science 2016, 352:712–716.

34. Castejon OJ, Fuller L, Dailey ME: Localization of synapsin-I and PSD-95 in developing postnatal rat cerebellar cortex. Brain Res Dev Brain Res 2004, 151:25–32.

35. Chi P, Greengard P, Ryan TA: Synapsin dispersion and reclustering during synaptic activity. Nat Neurosci 2001, 4:1187–1193.

36. De Camilli P, Cameron R, Greengard P: Synapsin I (protein I), a nerve terminal-specific phosphoprotein. I. Its general distribution in synapses of the central and peripheral nervous system demonstrated by immunofluorescence in frozen and plastic sections. J Cell Biol 1983, 96:1337–1354.

37. Nixon RA: The role of autophagy in neurodegenerative disease. Nat Med 2013, 19:983–997.

38. Nixon RA, Wegiel J, Kumar A, Yu WH, Peterhoff C, Cataldo A, Cuervo AM: Extensive involvement of autophagy in Alzheimer disease: an immuno-electron microscopy study. J Neuropathol Exp Neurol 2005, 64:113–122.

39. Kang R, Zeh HJ, Lotze MT, Tang D: The Beclin 1 network regulates autophagy and apoptosis. Cell Death Differ 2011, 18:571–580.

40. Tran S, Fairlie WD, Lee EF: BECLIN1: Protein Structure, Function and Regulation. Cells 2021, 10.

41. Tanida I, Ueno T, Kominami E: LC3 and Autophagy. Methods Mol Biol 2008, 445:77–88.

42. Fan FC, Liu LM, Guo M, Du Y, Chen YW, Loh YP, Cheng Y: Neurotrophic factor-alpha1/carboxypeptidase E controls progression and reversal of Alzheimer’s disease pathogenesis in mice. Theranostics 2025, 15:2279–2292.

43. Jiang D, Liu H, Li T, Zhao S, Yang K, Yao F, Zhou B, Feng H, Wang S, Shen J, et al: Agomirs upregulating carboxypeptidase E expression rescue hippocampal neurogenesis and memory deficits in Alzheimer’s disease. Transl Neurodegener 2024, 13:24.

44. Chun Y, Kim J: Autophagy: An Essential Degradation Program for Cellular Homeostasis and Life. Cells 2018, 7.

45. Rousseaux MW, de Haro M, Lasagna-Reeves CA, De Maio A, Park J, Jafar-Nejad P, Al-Ramahi I, Sharma A, See L, Lu N, et al: TRIM28 regulates the nuclear accumulation and toxicity of both alpha-synuclein and tau. Elife 2016, 5.

46. Rousseaux MW, Revelli JP, Vazquez-Velez GE, Kim JY, Craigen E, Gonzales K, Beckinghausen J, Zoghbi HY: Depleting Trim28 in adult mice is well tolerated and reduces levels of alpha-synuclein and tau. Elife 2018, 7.

47. Kim NY, Cho MH, Won SH, Kang HJ, Yoon SY, Kim DH: Sorting nexin-4 regulates beta-amyloid production by modulating beta-site-activating cleavage enzyme-1. Alzheimers Res Ther 2017, 9:4.

48. Pichet Binette A, Gaiteri C, Wennstrom M, Kumar A, Hristovska I, Spotorno N, Salvado G, Strandberg O, Mathys H, Tsai LH, et al: Proteomic changes in Alzheimer’s disease associated with progressive Abeta plaque and tau tangle pathologies. Nat Neurosci 2024, 27:1880–1891.

49. Ali M, Timsina J, Western D, Liu M, Beric A, Budde J, Do A, Heo G, Wang L, Gentsch J, et al: Multi-cohort cerebrospinal fluid proteomics identifies robust molecular signatures across the Alzheimer disease continuum. Neuron 2025, 113:1363–1379 e1369.

50. Yarbro JM, Han X, Dasgupta A, Yang K, Liu D, Shrestha HK, Zaman M, Wang Z, Yu K, Lee DG, et al: Human-mouse proteomics reveals the shared pathways in Alzheimer’s disease and delayed protein turnover in the amyloidome. bioRxiv 2024.

51. Askenazi M, Kavanagh T, Pires G, Ueberheide B, Wisniewski T, Drummond E: Compilation of reported protein changes in the brain in Alzheimer’s disease. Nat Commun 2023, 14:4466.

52. Pickford F, Masliah E, Britschgi M, Lucin K, Narasimhan R, Jaeger PA, Small S, Spencer B, Rockenstein E, Levine B, Wyss-Coray T: The autophagy-related protein beclin 1 shows reduced expression in early Alzheimer disease and regulates amyloid beta accumulation in mice. J Clin Invest 2008, 118:2190–2199.

53. Yang F, Beltran-Lobo P, Sung K, Goldrick C, Croft CL, Nishimura A, Hedges E, Mahiddine F, Troakes C, Golde TE, et al: Reactive astrocytes secrete the chaperone HSPB1 to mediate neuroprotection. Sci Adv 2024, 10:eadk9884.

54. van Dyck CH, Swanson CJ, Aisen P, Bateman RJ, Chen C, Gee M, Kanekiyo M, Li D, Reyderman L, Cohen S, et al: Lecanemab in Early Alzheimer’s Disease. N Engl J Med 2023, 388:9–21.

55. Sims JR, Zimmer JA, Evans CD, Lu M, Ardayfio P, Sparks J, Wessels AM, Shcherbinin S, Wang H, Monkul Nery ES, et al: Donanemab in Early Symptomatic Alzheimer Disease: The TRAILBLAZER-ALZ 2 Randomized Clinical Trial. JAMA 2023, 330:512–527.

56. Pickhardt M, Gazova Z, von Bergen M, Khlistunova I, Wang Y, Hascher A, Mandelkow EM, Biernat J, Mandelkow E: Anthraquinones inhibit tau aggregation and dissolve Alzheimer’s paired helical filaments in vitro and in cells. J Biol Chem 2005, 280:3628–3635.

57. Congdon EE, Ji C, Tetlow AM, Jiang Y, Sigurdsson EM: Tau-targeting therapies for Alzheimer disease: current status and future directions. Nat Rev Neurol 2023, 19:715–736.

58. Esquer A, Blanc F, Collongues N: Immunotherapies Targeting Amyloid and Tau Protein in Alzheimer’s Disease: Should We Move Away from Diseases and Focus on Biological Targets? A Systematic Review and Expert Opinion. Neurol Ther 2023, 12:1883–1907.

